# Transcriptome signature of *E. coli* under a prolonged starvation environment

**DOI:** 10.1101/2020.05.11.085241

**Authors:** Sotaro Takano, Hiromi Takahashi, Yoshie Yama, Ryo Miyazaki, Saburo Tsuru

**Author notes:** Address correspondence Saburo Tsuru.

## Abstract

**Background:** “Non-growing” is a dominant life form of microorganisms in nature, where available nutrients and resources are extremely limited. However, the knowledge of the manner in which microorganisms resist nutrient deficiency is still rudimentary compared to those of the growing cells. In laboratory culture, *Escherichia coli* can survive for several years under starvation, denoted as long-term stationary phase (LSP), where a small fraction of the cells survive by recycling resources released from the starved nonviable cells and constitute a model system for understanding survival mechanisms under long-term starvation. Although the physiology by which viable cells in LSP adapt to long-term starvation is of great interest, their genome-wide response has not yet been fully understood.

**Results:** To understand the physiological state of viable cells in the LSP environment, we analyzed the transcriptional profiles of cells exposed to the supernatant of LSP culture. We found that high expression of transporter genes and low expression of biosynthesis genes are the primary responses of the cells in the LSP supernatant compared to growing cells, which display similar responses to cells entering the stationary phase from the exponential growth phase. We also revealed some specific transcriptional responses in the LSP supernatant, such as higher expression of stress-response genes and lower expression of translation-related genes, compared to other non-growing conditions. This suggests that cells in LSP are highly efficient in terms of cellular survival and maintenance functions under starvation conditions. We also found population-density-dependent gene expression profiles in LSP, which are also informative to understand the survival mechanism of bacterial population.

**Conclusion:** Our current comprehensive analysis of the transcriptome of *E. coli* cells provides an overview of the genome-wide response to the long-term starvation environment. We detected both common and distinctive responses in the primary transcriptional changes between the short- and long-term stationary phase cultures, which could provide clues to understand the possible molecular mechanisms underlying survivability in the starved environment.

## Background

The physiology of growing microorganisms has been quantitatively characterized based on stable and reproducible culture systems since the 1940s [1]. The quantitative approaches have steadily revealed fundamental principles in exponentially growing microorganisms, such as the growth rate’s strong dependence on available nutrients in the environment or the cellular macromolecular composition (e.g., ribosomes, DNA, and RNA) [2-4]. The expression profiles of genes responsible for these interrelations have also been characterized [5-7], enabling us to understand the physiology of growing cells from systems-level mechanisms. However, most prokaryotes in nature live in nutrient-poor environments, such as deep biospheres, where they exhibit slow- or non-growing states [8]. The physiology of such microbial cells has been appreciated as a general topic [9,10], yet our understanding of how they survive in nutrient-poor conditions that provide marginal energy for cellular growth or the maintenance of basic cellular functions has remained rudimentary. To systematically understand the cellular physiology of starved microbes, key biological processes for their survival should be characterized based on quantitative measurements.

Microorganisms usually retain viability (reproductivity) during long-term periods of starvation [8-10]. In a laboratory culture system, *Escherichia coli* maintains viability for years after the exhaustion of nutrients [11]. The cells initially grow exponentially in a fresh nutrient medium, but growth is stopped after the consumption of nutrients, where cells enter short-term stationary phase. A majority of the cells die after the short-term stationary phase, but a small fraction of cells (0.1∼0.1 %) can continue surviving for months or years, which is termed as long-term stationary phase (LSP) [11]. Especially in the case of carbon starvation in synthetic minimal medium, the survival kinetics and rates of starved *E. coli* cells are quite reproducible, which enables an understanding of the long-term survival mechanisms based on quantitative measurements [12-14]. One important finding from these studies on carbon-starved *E. coli* cells is that cellular death and growth are strongly affected by nutrients released from dead cells [12,13]. A theoretical study also demonstrated that the constant survivability in LSP is accomplished by recycling dead cells and supported by its regulation of population density such that they recover their viability when viable cells decrease in the LSP [12].

Although studies have demonstrated cell-level behaviors responsible for constant survival in LSP [12,13], molecular functions or biological processes required for the maintenance of viability are still unknown, which impedes further understanding of the long-term survival mechanism regarding the physiological state of viable cells. The viable population in LSP can utilize chemicals released from starved cells but grow at a very slow rate [12]. Thus, it could be expected that growth-related cellular functions in the viable population must be activated by the nutrients from dead cells, but their physiological state must be different from those of the exponentially growing cells. Direct quantification of the transcriptome in long-term surviving cells would help elucidate the global transcriptional response occurring in prolonged starvation, but it is quite difficult to directly analyze such a small group of viable cells (0.1∼0.01 %) apart from a vast majority of the dead (or nonviable) population in LSP culture. Moreover, little is known about the genome-wide cellular processes specifically observed in the LSP survivors. The extraction of sufficient amounts of molecules from such a small number of cells is another technical challenge for any omics approach.

Given that survival in LSP relies on nutrients released from dead cells to the culture [12,13], the physiology of the survivors can be inferred by investigating the cellular response to the supernatant of LSP culture. Therefore, in this study, we analyzed the transcription of over 3,700 genes of *E. coli* exposed to the supernatant of LSP to forecast genome-wide responses under a prolonged starvation environment. Here, we used the LSP culture in carbon starvation as a model system to study the cellular response in the prolonged starvation environment. We compared the transcription profiles of cells in the LSP supernatant to those in the exponential growth phase, short-term stationary phase, or fresh glucose-deficient minimal medium and found distinct gene expression patterns in each condition. We also identified population-density-dependent transcriptome changes in cells exposed to the LSP supernatant, which supports the density-dependent physiological changes: fast death and slow growth at high cell density in LSP, from molecular-level mechanisms.

## Results

### Experimental design

The experimental design for investigating gene expression profiles in four different conditions is shown in Fig. 1A. In each condition, *E. coli* was first grown in a glucose-rich synthetic medium (M63 medium with 22.2 mM glucose; for details, see Materials and Methods) for 20 h until late-log phase (∼10^9^ cells/mL). We designated this cell culture as GLC+ (late-log). We divided the GLC+ (late-log) culture into two aliquots and further incubated one for 16 h (then, this culture reached the plateau (Additional file 1: Fig. S1, inset)) and designated it as GLC+ (ST). We washed the other aliquot, resuspended it with the LSP supernatant (i.e., a supernatant of the *E. coli* culture in M63 medium without glucose for 30 days) or a glucose-deficient medium (i.e., fresh M63 medium without glucose), and incubated it for 16 h. These two cell cultures were designated as LSPE (cells cultured in <u>L</u>ong-term <u>S</u>tationary Phase <u>E</u>nvironment) or GLC−. As the incubation time increased, the viability of these cell cultures gradually decreased in all conditions (Additional file 1: Fig. S1), resulting in an increase in the number of dead cells. We focused on the transcriptional response specific to a viable population; thus, contamination of dead cells in the cultures was not ideal for the transcriptome analysis. The results of the viability assays revealed that 16 h of incubation did not reduce the survival rate in all culture conditions (Additional file 1: Fig. S1); thus, we collected samples from all the cell cultures for RNA isolation after 16 h from transferring to each condition. In addition, a 16-h incubation period in the GLC+ medium was sufficient to see the entry into the short-term stationary phase, where the growth rate reached almost 0 (Additional file 1: Fig. S1, inset). Thus, collecting the samples at this time point should be sufficient for analyzing the physiological differences between the GLC+ (late-log) and GLC+ (ST) samples. We also found that LSPE showed different survival kinetics from GLC−; while the survival rate gradually decreased over time in both cultures (Additional file 1: Fig. S1), the death rate calculated from the survival curves was significantly lower in LSPE (p-value < 0.05, Fig. 1B). Therefore, culturing the cells in the LSP supernatant attenuated a decrease in the survival rate compared to that of the cells in GLC−.

**Fig. 1.**
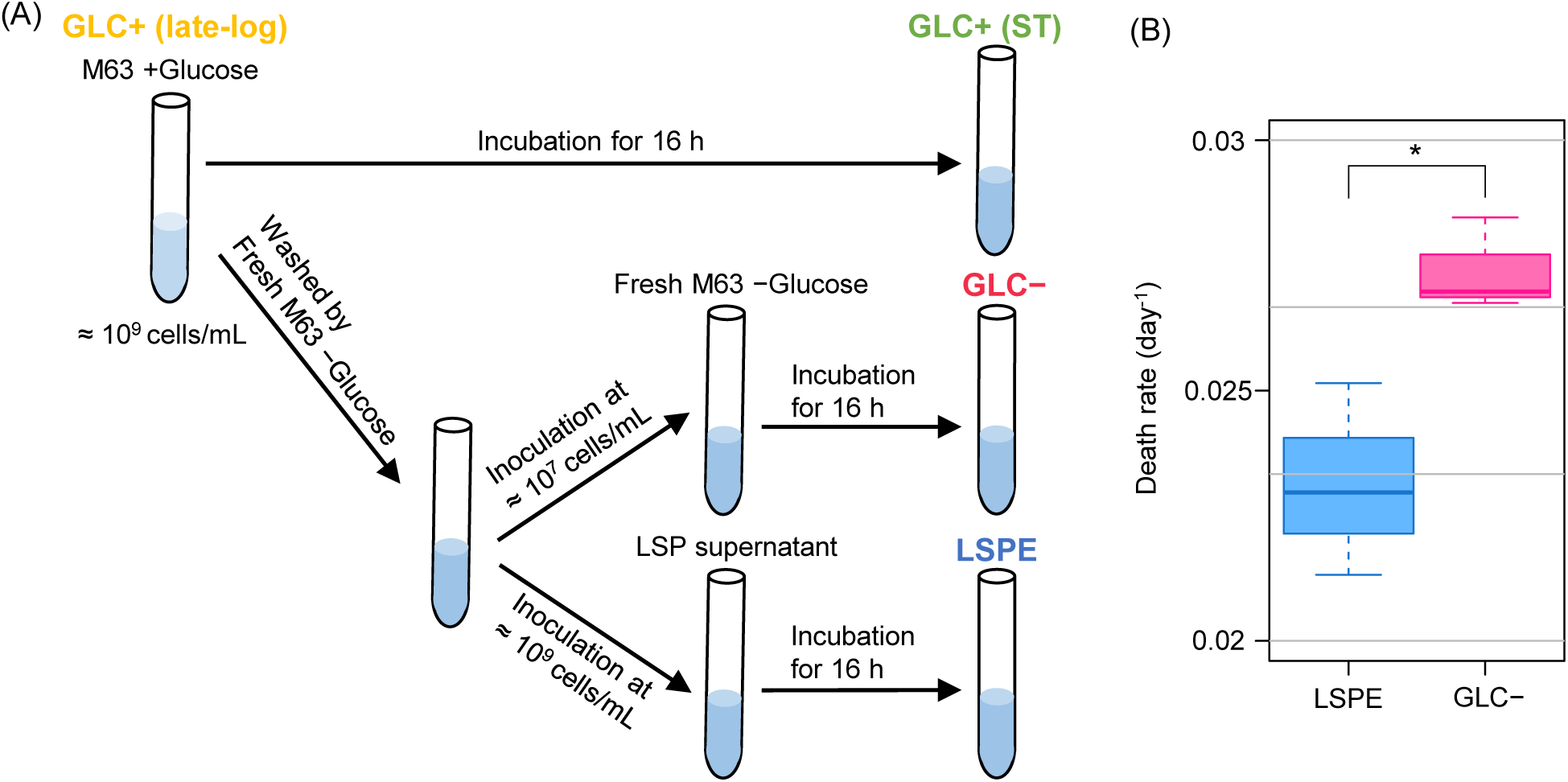
Experimental design for the transcriptome analysis. (A) We first cultured *Escherichia coli* cells in M63 minimal medium with glucose and collected them from the pre-grown culture (labeled as GLC+ (late-log)). Then, this culture was divided into two aliquots. One divided aliquot from the late-log phase culture was incubated for 16 h to prepare the short-term stationary phase cell culture (labeled as GLC+ (ST)). To prepare the LSPE or GLC− cultures, we washed the other aliquot with fresh M63 minimal medium without glucose and inoculated it into the LSP supernatant or fresh M63 minimal medium (without glucose). Given that incubation of the cells changes the culture condition by the release of molecules from the cells during starvation [12], we inoculated the cells at 10^7^ cells/mL for the analysis in the case of GLC− to attenuate the effect of molecules released during experiments. (B) Comparison of death rates between cells in the LSP supernatant (LSPE) and fresh glucose-deficient medium (GLC−). Death rates were estimated by the fitting of the survival curve (Additional file 1: Fig. S1) to a single exponential equation (see Materials and Methods). Error bars indicate the standard deviation of biological replicates (n = 3). Asterisks indicate statistical significance levels of Welch’s two-sample *t*-test (**, p < 0.01; *, p < 0.05).

### Expression profiles in LSPE, GLC+ (ST), and GLC− are similarly grouped together

We first performed hierarchical clustering for all analyzed samples across all genes to compare gene expression profiles. We found that GLC+ (late-log) showed different gene expression profiles from the other three conditions (hereafter the three conditions were called “non-growing group”) (Fig. 2A). In the “non-growing group,” GLC− was further separated from the other two conditions. Principal component analysis (PCA) suggests a similar tendency in the transcriptional profile: LSPE and GLC+ (ST) are clustered, whereas GLC+ (late-log) and GLC− are distinct (Fig. 2B). These results clearly indicate that, while transcriptomes between the growing and non-growing conditions are primarily different, there are distinct characteristics within the “non-growing group.” As LSPE showed a more similar gene expression profile to GLC+ (ST) than GLC−, we further grouped LSPE and GLC+ (ST) together and designated this group as SLSP (<u>s</u>hort- and long-term <u>s</u>tationary <u>p</u>hase).

**Fig. 2.**
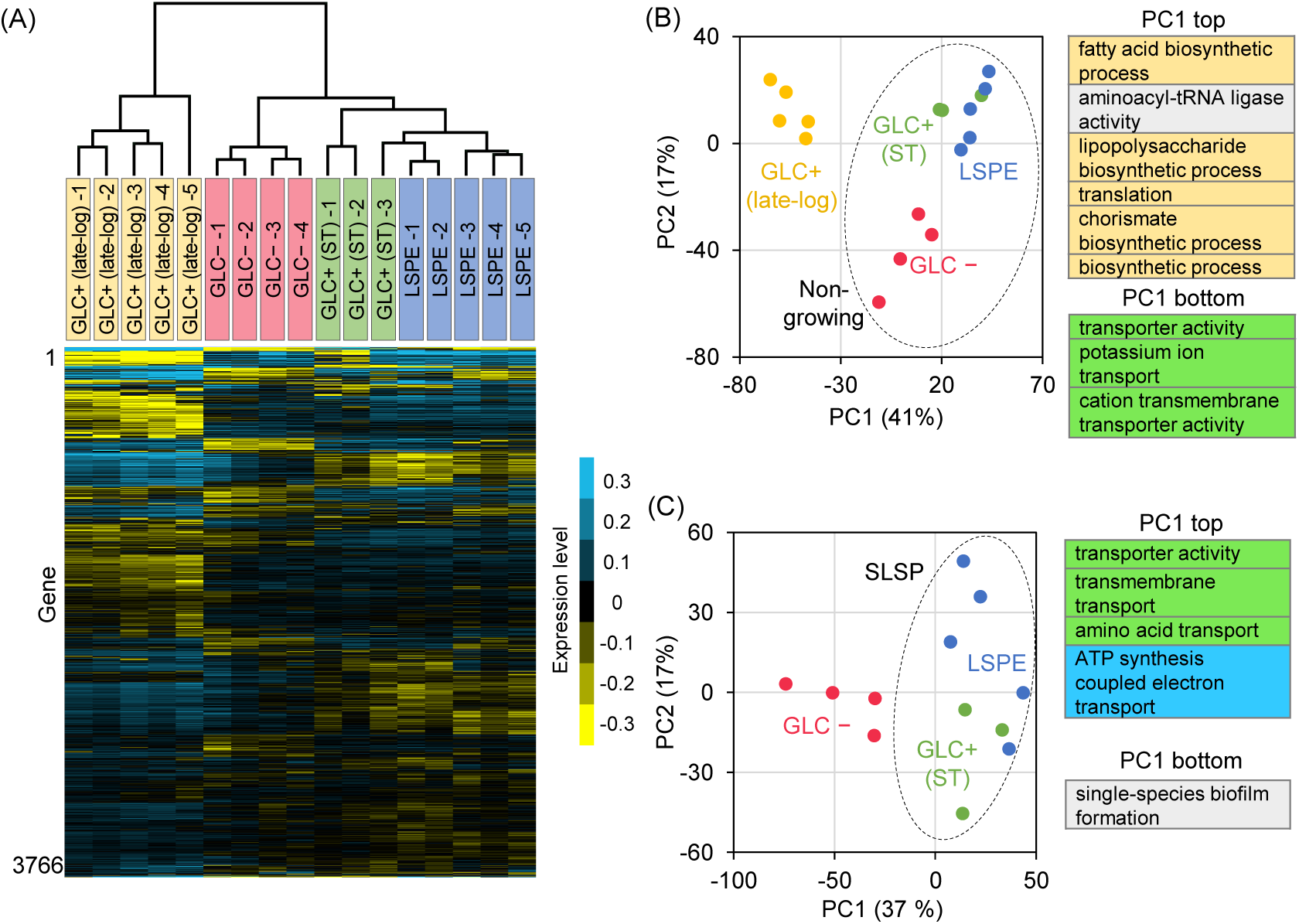
Comprehensive analysis of gene expression data. (A) Heatmap of the expression levels in 3,766 genes in the LSP supernatant (LSPE, n = 5), the fresh glucose-deficient culture (GLC−, n = 4), the late-log phase culture in the presence of glucose (GLC+ (late-log), n = 5), or the short-term stationary phase culture (GLC+ (ST), n = 3). In each gene, the expression data were normalized to the mean value of all samples. We performed clustering both by genes and arrays using the average linkage method based on the Euclidean distance among the samples or genes. (B) Plots of PCA scores from the analysis using all analyzed samples. Plots indicate scores of individual samples in each condition. The percentage displayed on each axis is the rate of the contribution of PC1 or PC2. We circled the non-growing group using a black dotted line. (C) Plots of PCA scores from the analysis within the non-growing group. We circled SLSP using a black dotted line. PC1 top or bottom groups are GO terms that have significant positive or negative contributions to PC1. GO terms associated with three functions, “transport,” “biosynthetic process,” and “oxidative phosphorylation” are colored green, yellow, and blue. All extracted gene sets are shown in Data S2 (Additional file 6).

### Screening of the differentially expressed gene sets between growing and non-growing groups

Given that LSPE was grouped into the non-growing group, we first characterized functional gene sets that were differentially expressed in the non-growing group compared to GLC+ (late-log). *K*-means clustering (*K =* 2) indicates that PC1 in Fig. 2B can explain the difference between those two groups (Additional file 2: Fig. S2A). Additionally, we statistically screened functional gene sets in which highly contributing genes to PC1 were significantly enriched. We used GO terms for categorizing genes by their functions [15] (Additional file 5: Data S1). We here focus on the difference in the cellular processes; thus, gene sets classified by their biological and molecular functions (i.e., child classes or parts of “biological processes (GO:0008150)” and “molecular functions (GO:0003674)”) were shown (Fig. 2B). There are two types of screened gene sets: PC1 top and PC1 bottom (Additional file 3: Fig. S3). Gene sets screened as “PC1 top” include genes positively correlated to PC1 scores in their expression levels (Additional file 3: Fig. S3). For instance, *caiT*, a member of “transporter activity (GO:0005215)” in the “PC1 top” group, showed a strong positive correlation with the PC1 scores (Additional file 3: Fig. S3), and the level of expression of this gene was higher in the non-growing group (Additional file 3: Fig. S3). In contrast, the expression levels in the “PC1 bottom” group were negatively correlated with the PC1 scores. *rpmF*, which is included in “translation (GO:0005840)” in the “PC1 bottom” group, showed strong negative loading for PC1 (Additional file 3: Fig. S3), and GLC+ (late-log) samples showed higher expression levels than the non-growing group (Additional file 3: Fig. S3).

The “PC1 top” gene sets were mostly associated with transport activity (i.e., child classes or parts of the “transporter activity (GO:0005215)” or “transport (GO:0006810)”), whereas the “PC1 bottom” were occupied by biosynthetic gene sets (i.e., child classes or parts of the “biosynthetic process (GO:0009058)”). Therefore, higher expression levels of transport genes and lower expression levels of biosynthesis genes would be major transcriptional changes in the non-growing group compared to GLC+ (late-log). This tendency was demonstrated by comparing the average transcripts levels of all gene members in “biosynthetic process (GO:0009058)” and “transport (GO:0006810)” (Additional file 4: Fig. S4). These functional categories are upper level GO terms associated with transport and biosynthesis function and comprise over 200 child classes (see Additional file 7: Data S3). As expected from the results of PCA, the lower expression levels in the “biosynthetic process (GO:0009058)” genes are common transcriptional changes among the non-growing group (Additional file 4: Fig. S4, GLC+ (ST), LSPE, and GLC−). In contrast, the increase in the gene expression of “transport (GO:0006810)” was specifically observed in SLSP (Additional file 4: Fig. S4, GLC+ (ST) and LSPE), indicating that high expression levels of transporter genes would be unique to the SLSP group.

### Screening of differentially expressed gene categories within the non-growing group

To investigate the difference in physiology between SLSP and GLC−, we next characterized the different expression profiles within the non-growing group. The result of *K*-means clustering indicated that PC1 in Fig. 2B cannot explain the difference between SLSP and GLC−(*K* = 3, 4, Additional file 2: Fig. S2A). Then, we performed PCA within the non-growing group (Fig. 2C). *K*-means clustering (*K* = 2) of the samples in the non-growing group by PC1 in Fig. 2C indicated that the SLSP group are distinguished from GLC− by the first principal component (Additional file 2: Fig. S2B); then, we characterized the differentially expressed gene categories by PC1 in Fig. 2C. The “PC1 top” group mostly associated with transport activity (e.g. “transporter activity (GO:0005215)” and “transmembrane transport (GO:0055085)”). As shown in Fig. S4 (Additional file 4), the higher expression levels in “transporter activity (GO:0005215)” were specifically observed in LSPE and GLC+ (ST). Thus, this characteristic is unique to SLSP. Another screened gene set related to oxidative phosphorylation (e.g. “ATP synthesis coupled electron transport (GO:0042773)”) (Fig. 2C) shows higher expression levels on average in SLSP than GLC− (Additional file 4: Fig. S4). In contrast, “single-species biofilm formation (GO:0044010)” was screened as the “PC1 bottom” category. The members showing low factor loading in this gene set are dinJ-yafQ and yoeB-yefM (Additional file 6: Data S2), both of which are the genes of toxin-antitoxin systems and are responsible for the regulation of RNA degradation [16].

### Comparison of expression profiles between in LSPE and GLC+ (ST)

To understand cellular responses specific to long-term, but not short-term, stationary phase environment, we identified the functional gene sets (GO terms) differentially expressed in LSPE and GLC+ (ST) by Gene Set Enrichment Analysis (GSEA) [17]. Significantly highly expressed genes in LSPE were genes for “phage shock (GO:0009271),” one member of “stress response” (Fig. 3A), which are known to maintain membrane integrity under stress conditions to avoid leakage of cytoplasmic contents and the loss of proton motive force [18-20]. Another highly expressed functional category is “transport” such as “transmembrane transport (GO:0055085)” in LSPE. Especially, “carbohydrate transport (GO:0008643)” shows high enrichment scores in the group of “transport” (Fig. 3A), indicating that LSPE samples would more actively take in ambient carbohydrates by increasing the expression levels of those gene sets. In contrast, gene expression related to “biosynthesis”, such as “translation (GO:0006412),” and catabolism & energy production, such as “aerobic respiration (GO:0009060)” and “tricarboxylic acid (TCA) cycle (GO:0006099),” were lower in LSPE than in GLC+ (ST) (Fig. 3A). These gene sets are associated with protein production and ATP concentration, which are strongly related to cellular growth [5, 21]. These results suggest that cells in the short-term stationary phase (GLC+ (ST)) somewhat retain their characteristic physiology in the growth phase, whereas LSPE cells exhibit more static and stress/survival-weighed physiology. To explore the balance between growth- and stress/survival-related cellular functions in LSPE cells, we compared the average expression levels of these two gene sets among LSPE, GLC+ (ST), and GLC+ (late-log). We used GO categories of “macromolecule biosynthetic process (MBS)” (GO:0009059) as a gene set for cell growth, because it includes a large number of growth-related genes [22], and “response to stress (RS)” (GO:0006950) as a gene set for stress survival (Additional file 7: Data S3), and then calculated a ratio of their average expression levels (AVR_RS_ / AVR_MBS_). The ratio was significantly higher in LSPE than in GLC+ (late-log) (p-value < 0.01, Fig. 3B), suggesting that the cells in the LSP environment weighted stress/survival tasks. In contrast, the ratio was lower in GLC+ (ST) than in LSPE, and there was no significant difference between GLC+ (ST) and GLC (late-log), supporting that LSPE would be in a more stress/survival-weighed physiology than GLC+ (ST).

**Fig. 3.**
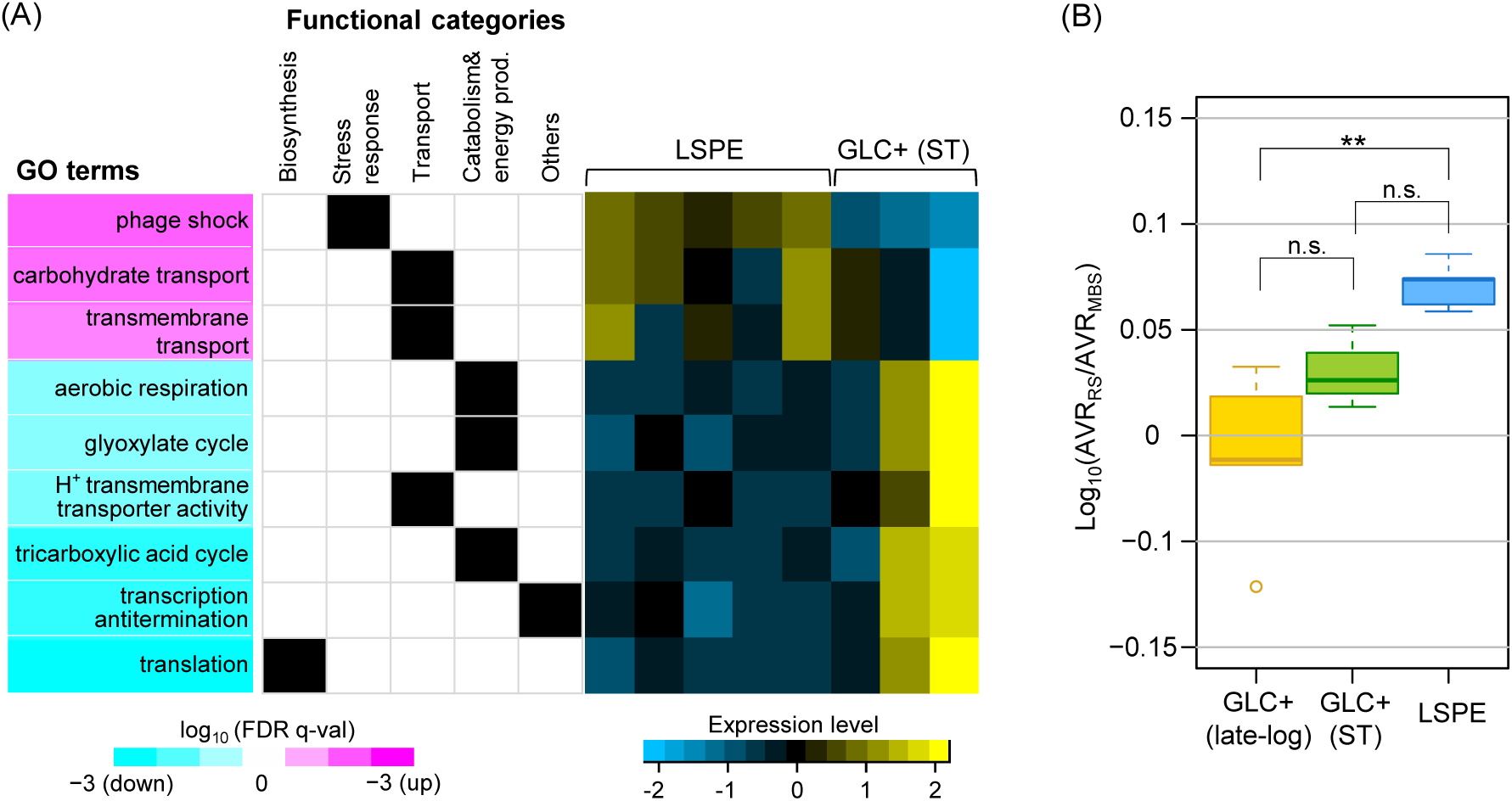
Differentially expressed gene categories in LSPE compared to GLC+ (ST). (A) Representative differentially expressed gene categories extracted by GSEA (false Discovery Rate (FDR), q-value < 0.05, for details, see Material and Methods). Statistical significance levels (FDR, q-value) are shown as heatmaps. Magenta in a heat map means that those gene sets were more highly expressed in LSPE, whereas cyan means that the expression levels of those gene sets are higher in GLC+ (ST). These screened gene sets were further grouped into five large categories: 1) biosynthesis (child classes or parts of “biosynthetic process (GO:0009058)”), 2) stress response (child classes or parts of “response to stress (GO:0006950)” and “anti-oxidant activity (GO:0016209)”), 3) transport (child classes and parts of “transport (GO:0006810)” or “transporter activity (GO:0005215)”), 4) catabolism and energy production (child classes and parts of “generation of precursor metabolites and energy” (GO:0006091) or “cellular catabolic process” (GO:0044248)), and 5) other categories. If each GO term was grouped into an indicated functional category, the box was filled with black. A yellow-blue scaled heatmap shows the expression levels in each GO term (i.e., the average gene expression levels of all members in each GO term) in biological replicates (LSPE: n = 5; GLC+ (ST): n = 3). The values of mRNA signals in each extracted GO category were normalized so that the average and the standard deviation in all replicates are 0 and 1. FDR q-values of all extracted categories are shown in Data S4 (Additional file 8). (B) log10 scaled ratio between the average mRNA signals of all members in “macromolecule biosynthetic process (GO:0009059),” designated as “MBS,” and “response to stress (GO:0006950),” designated as “RS.” Error bars indicate the standard deviation of average signals among the biological replicates. Asterisks indicate statistical significance levels according to Tukey’s HSD test (**, p < 0.01; *, p < 0.05).

### Density-dependent transcriptional changes in the LSP supernatant

Previous studies demonstrated that cellular growth and death in the spent medium under carbon starvation are affected by the density of viable cells [12,14]. Thus, we expected that cellular viability in LSPE would be also affected by the density of the cells. To test this possibility, we measured the viability of cells inoculated at different densities in the LSP supernatant. As expected, cellular viability in LSPE was significantly changed by inoculated cell density: the larger the density, the less the viability becomes (Fig. 4A). We then analyzed the transcriptomes of those cells in LSPE at different cell concentrations (10^7^ or 10^9^ cells/mL). Although a very strong positive correlation in gene expression was observed between those two conditions (Fig 4B), we also detected 54 differentially expressed gene sets by GSEA (see Additional file 9: Data S5 for all extracted gene sets). In the lower cell density (10^7^ cells/mL), genes for biosynthesis (i.e., “translation (GO:0006412)” and “fatty acid biosynthetic process (GO:0006633)”), transcription (i.e., “DNA-directed RNA polymerase activity (GO:0003899)”), and cell division (i.e., “barrier septum assembly (GO: GO:0000917)”) were significantly expressed (Fig. 4C). The levels of expression related to stress response and maintenance (i.e., “protein folding (GO:0006457)”, “anti-oxidant (GO:0016209)”, and “response to heat (GO:0009408)”) were also high (Fig. 4C). These suggest that cells in LSP at low density are more competent for reproduction, accompanying the expression of stress response genes, than those at higher density. In the higher density (10^9^ cells/mL), contrastingly, a large number of genes involved in transport functions were highly expressed (Fig. 4C). For instance, “amino acid transport (GO:0006865)”, “carbohydrate transport (GO:0008643)”, and “ion transport (GO:0006811)” were highly expressed transport genes (Fig. 4C), which are responsible for the uptake of extracellular molecules. A gene set of “NADH dehydrogenase (ubiquinone) activity” also showed a higher expression level, implying that cells in LSP at high density a more likely to commit to the oxidative phosphorylation process for ATP production than those at low cell density.

**Fig. 4.**
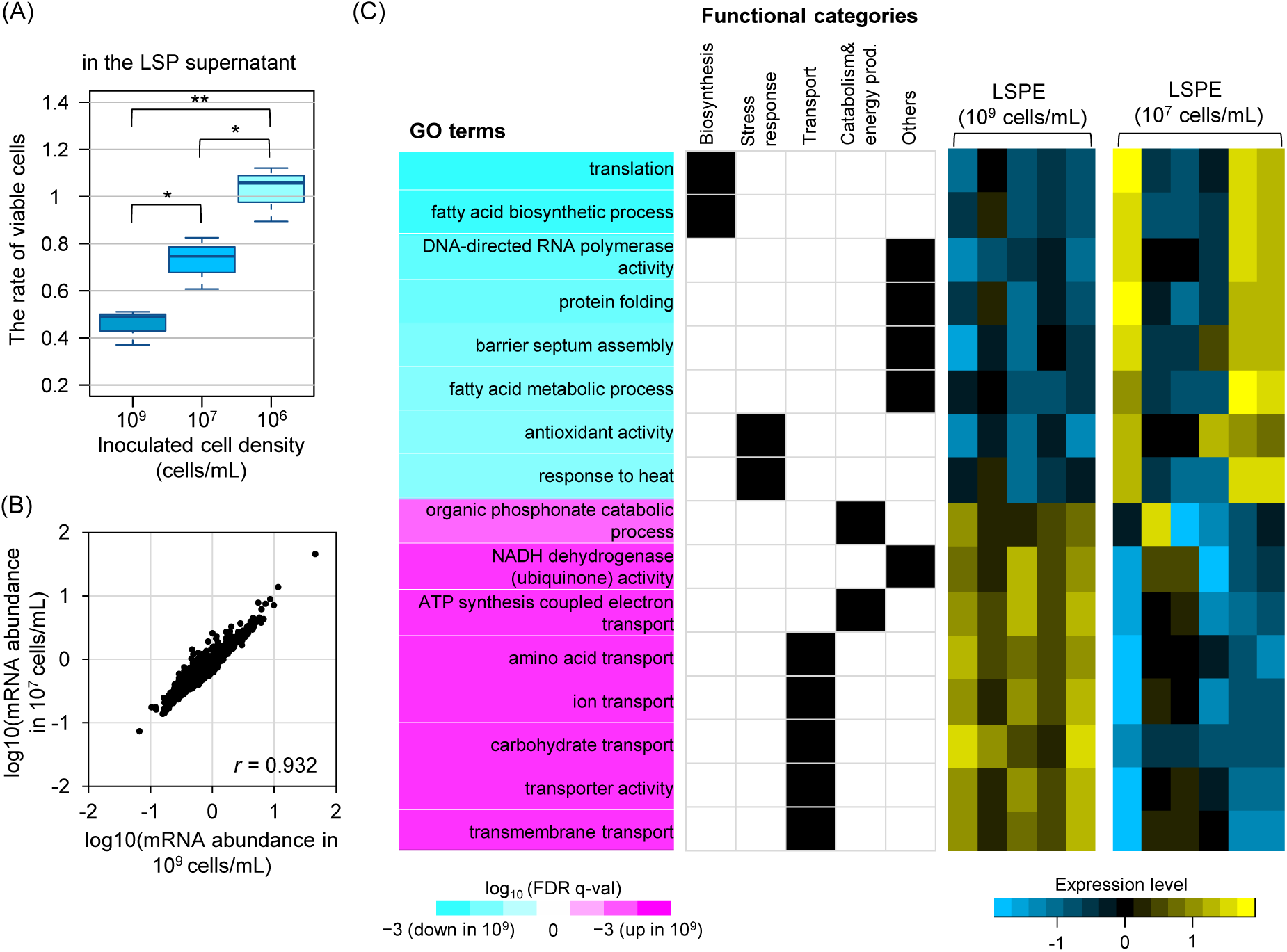
Differentially expressed genes depending on the population density in the LSP environment. (A) The changes in viability depending on the population density in the LSP supernatant. The rate of viable cells after 3 days’ incubation at different population densities is shown as boxplots. Error bars indicate standard deviations of biological triplicate experiments. Asterisks indicate statistical significance levels according to Tukey’s HSD test (**, p < 0.01; *, p < 0.05). (B) Dot plots of the expression levels in the condition of 10^9^ and 10^7^ cells/mL. *r* refers to Pearson correlation coefficient. (C) The result of GSEA comparing 10^7^ and 10^9^ cells/mL conditions is shown as heatmaps of FDR q-value. We compared expression levels at 10^9^ cells/mL to 10^7^ cells/mL, and gene sets showing higher expression levels in the condition of 10^9^ cells/mL are colored magenta, and gene sets whose expression levels are higher in 10^7^ cells/mL are colored cyan. We also categorized these enriched gene sets into five functional groups in the same manner as done in Fig. 3A. The statistical significance level is indicated by the brightness of color. The classification of the gene sets was done in the same manner as done in Fig. 3. We also showed the average expression levels of all members in each GO term in biological replicates (10^9^ cells/mL, n = 5; 10^7^ cells/mL, n = 6) with the same normalization method as in Fig. 3A.

## Discussion

### A common transcriptional response in non-growing condition

Our transcriptome analysis revealed a common transcriptional response in different starved cultures. Compared to cells in exponentially growing phase (GLC+ (late-log)), one of the major transcriptional changes in long-term stationary phase (LSP) is a decrease of gene expression for biosynthesis of fatty acids, lipopolysaccharides, proteins, and chorismates (i.e., precursors for aromatic amino acid compounds) (Fig. 2B and S4), which is similar to the expression profiles in other non-growing conditions, such as short-term stationary phase (GLC+ (ST)) or glucose-deficient medium (GLC−). As these gene sets are involved in the synthesis of major cellular components and essential for cellular growth in *E. coli* [23,24], their low expression levels would be one of the possible mechanisms to explain non-growing state of cells in LSP [14,15], which is a common tendency of bacterial transcriptional response after halting growth [25-29].

Among the non-growing conditions, we also revealed that the expression profiles of LSPE were more similar to those of GLC+ (ST) than GLC−. LSPE and GLC+ (ST) showed higher expression levels in genes associated with transport and oxidative phosphorylation than GLC−. Among all the tested conditions, the high expression levels in “transport (GO:0006810)” gene sets are unique to LSPE and GLC+ (ST) (Fig. 2 and S4). The cultures of LSPE and GLC+ (ST) must contain various chemical compounds released from cells (e.g., carbohydrates, nucleotides, and amino-acids) [30-32], and a high expression of various transporter genes to be beneficial for cells to import and recycle ambient nutrients for their survival. Furthermore, the importance of the ability to utilize compounds released from dead cells in the LSP supernatant has been previously reported [12,14,30]. Additionally, the scavenging of remaining nutrients is one of the major responses in the short-term stationary phase as previous studies suggested [33,34]. High expression of “ATP synthesis coupled electron transport (GO:0042773)” in LSPE and GLC+ (ST) would also link to those metabolic activities by facilitating ATP production from the imported molecules or producing energy for the production of the transporter proteins. Activation of transport functions and ATP synthesis would be one of the important mechanisms for constant survival in short- and long-term stationary phases.

### Transcriptional responses specific to the long-term stationary phase environment

Although LSPE has similar transcriptional characteristics to GLC+ (ST), several functional gene sets were differentially expressed between populations. One of the major differences is higher expression levels of protein-synthesis genes (e.g., translation) in GLC+ (ST) than in LSPE. Protein synthesis is an energy-consuming process tightly coupled to cellular growth in bacteria [3,4,35]. Therefore, the higher expression levels of those gene sets indicate more growth-weighed physiology in short-term stationary phase than LSP. A previous study also demonstrated that *E. coli* cells in short-term stationary phase can constantly produce proteins after the stop of growth [36], suggesting that those cells still partially inherit their physiology during the growing phase. Conversely, phage shock protein family, one of the important cell envelope stress response genes, was strongly induced in LSPE (Fig. 3A), suggesting that the physiology of LSP would be a more stress-resistant state rather than a reproductive state. We speculate that this stress/survival-weighed transcriptional change would contribute to survival in LSPE. In addition, a previous study reported that phage shock protein defective mutants exhibited less survivability in a certain condition of stationary phase culture [37]. Moreover, induction of phage shock proteins seems to lead to repression of aerobic respiration and increase of anaerobic respiration and fermentation [38] through ArcA/ArcB system, suggesting downregulation of energy production for fast growth.

### Density-dependent gene expressions in the LSP condition

Another discovery of this study is the density-dependent gene expression in the LSP supernatant. Inoculation of cells at the lower cell density in the LSP supernatant (10^7^ cells/mL) enhanced the survivability of the cells with higher expressions of genes for protein production, cell division, and cellular maintenance (Fig. 4). On the other hand, the cells at high density (10^9^ cells/mL) reduce those transcripts but increase the expression of energy metabolism and transporter genes (Fig. 4C). High expression levels of energy metabolism genes would facilitate ATP production and the maintenance of high-energetic-cost processes such as production of transporters to increase affinity to ambient nutrients [39]. Bacterial cells generally increase the expression of genes related to uptake of nutrients when the concentrations of these ambient chemicals decreased [40-42]. The amount of extracellular resources per cell is deemed to be small in the case of high cell density, and thus the expression of genes encoding transporter and energy metabolism would be advantageous in the condition of high cell density. In contrast, when the viable cell density is low, the amount of substrates per cell is relatively high, and the devotion of many resources to transcription of transporter and energy metabolism genes would be not beneficial. Instead, allocating their energy and resources for the cell division and maintenance would be more advantageous for preventing a decrease of a viable population in LSP.

## Conclusions

Previous studies demonstrated the survival mechanisms and role of recycling activity in LSP by mainly quantitating cell-level physiological traits (e.g., growth and death rate) using carbon-starved *E.coli* cells; however, the underlying molecular mechanisms and biological processes for the long-term survival have still not yet been cleared. Our study using DNA microarray characterized the genome-wide response of cells in the environment of LSP and screened potentially important cellular processes for long-term survival. Besides, our transcriptome analysis revealed transcriptional changes by population density in LSP, which are also informative to understand the density-dependent changes in cell-level behaviors from the view of molecular functions. Although a detailed gene regulatory network for realizing the cellular physiology in the LSP environment is still unclear, our results propose candidates for such important gene members and pave the way to understand survival mechanisms during prolonged starvation from molecular-level systems.

## Methods

### Strain and media

We used the *E. coli* strain MDS42, which was purchased from Scarab Genomics (Madison, WI, USA), for all experiments. The bacterial cells were cultured with M63 minimal medium (62 mM K_2_HPO_4_, 39 mM KH_2_PO_4_, 15 mM (NH_4_)_2_SO_4_, 2 µM FeSO_4_ · 7H_2_O, 200 µM MgSO_4_ · 7H_2_O), and 22 mM glucose was supplemented as necessary.

### Culture conditions and RNA sample preparation for DNA microarray

The *E. coli* cells were inoculated into fresh M63 medium with glucose from glycerol stock and grown for 20 h at a density of approximately 1 × 10^9^ cells/mL. Cells were then shaken at 160 rpm in a BR-21FP air incubator (Taitec, Saitama, Japan) at 37°C aerobically. We divided this culture into two aliquots. One aliquot of the grown cultures further incubated at 37 °C for 16 h to analyze transcriptome in short-term stationary phase culture conditions (GLC+ (ST)). For the transcriptome analysis in the GLC− or LSPE, the other aliquot was centrifuged at 7500 × g for 3 min and resuspended by M63 minimal medium without glucose. These washing steps were performed three times to eliminate glucose in the culture, and then samples were inoculated into each condition (see also Fig. 1A). To cultivate a sufficient amount of mRNA molecules from the culture, we cultured cells at a concentration of > 10^7^ cells/mL in all experimental conditions and collected > 10^9^ cells from each sample. To harvest the samples for GeneChip analysis, the cell cultures were mixed into chilled phenol-ethanol solution (1 g of phenol in 10 mL of ethanol) and centrifuged at 7500 × g for 3 min at 4 °C. After eliminating the supernatants, the cell pellets were frozen by liquid nitrogen and stored at −80 °C. Total RNA was extracted using a PureLink Mini kit (Thermo Fisher Scientific, Waltham, MA, USA) according to the manufacturer’s protocol.

### Microarrays and data normalization

The cDNA synthesis from the purified RNA, fragmentation, labeling, and hybridization of cDNA were performed in accordance with the manufacturer’s protocol of Affymetrix (Santa Clara, CA, USA). A high-density DNA microarray that covers the whole genome of *E. coli* W3110 strain was used for analysis. Microarray data were extracted based on the finite hybridization model as previously described [43,44], and the data sets of genes in which average values were less than −1.5 pM were eliminated to prevent the effects of noise derived from the small values as described previously [45]. These data sets were further normalized so that all data sets have common statistics on a logarithmic scale [46]. Finally, we used a total of 3,766 genes for the following analysis.

### Computational data analysis

Clustering of genes and arrays according to 3,766 transcriptional profiles was performed by Cluster 3.0 [47] using the average linkage method of the Euclidean distance.

A GO term annotation was used for functional classification of genes in all transcriptome analysis, and the annotation list of *E. coli* strain K-12 was collected from Ecocyc [15] (http://www.biocyc.org), with slight modifications (Additional file 5: Data S1). To make 3766-dimensional data to low dimension and visualize simply, we performed PCA and *K-*means clustering employing R-package [48] using the correlation matrices of the normalized data sets. If factor loading is i) in the top or bottom 3% among all analyzed genes, and ii) more than 0.8 or less than −0.8, we identified those genes as highly contributing genes to each PC score. For screening GO terms including a significantly large number of genes with high loading-factor, we used a hypergeometric distribution (p < 0.01) as previously described [49]. For studying differentially expressed gene sets, we conducted Gene Set Enrichment Analysis (GSEA) [17]. We excluded GO terms consisting of less than 6 genes or more than 200 genes for GSEA, PCA, and post hoc analysis.

### Viability assay

The number of viable cells was estimated by colony-forming unit (CFU) in all experiments. Cell cultures incubated in each condition were harvested, diluted, and inoculated onto M63 agar (1.5%) plates supplemented with 0.4 % glucose. The duplicate inoculated plates were incubated at 37 °C for approximately 2 d, and the number of colonies was manually counted.

### Estimation of relative death rate using multiple time points on survival curves

In Fig. S1 (Additional file 1), the relative death rate *µ* was estimated by the fitting to a following single exponential equation.

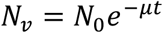

Here, *N*_*v*_ and *N*_*0*_ denote the number of viable cells and initial cell density in the culture. We used polyfit function in MATLAB (MathWorks, Natick, MA, USA) for fitting to the experimental data.

### Preparation of supernatants from LSP cultures

We grew *E. coli* cells in M63 with glucose and washed the cells thrice with M63 without glucose, as described above. The glucose-starved cultures were incubated for 30 d at 37 °C aerobically with shaking at 160 rpm. After a 30 d incubation, the cultures were harvested by centrifugation at 7500 × g for 3 min at room temperature. The supernatants were filtered with a 0.22-µm filter (EMD Millipore, Billerica, MA, USA) and stored at 4 °C.

## Declarations

### Ethics approval and consent to participate

Not applicable.

### Consent for publication

Not applicable.

### Availability of data and materials

The raw CEL files used for the all transcriptome analysis here were deposited in the NCBI Gene Expression Omnibus database (accession no. GSE149236).

### Competing interests

The authors declare that they have no competing interests.

### Funding

This work was partly supported by Japan Society for the Promotion of Science KAKENHI Grants 18H02427, 15KT0078 for SBT, and Grant-in-Aid for JSPS Fellows 14J02139 for ST. The funders had no role in study design, data collection, and interpretation or the decision to submit the work for publication.

### Author contributions

ST and SBT designed research; ST, YY, and HT performed experiments and analyzed data; and ST, RM, and SBT wrote the paper.

## Acknowledgments

We thank members of the Symbiotic Network Design Laboratory (Osaka University) and Microbial and Genetic Resources Research Group (AIST) for discussions and valuable comments.

## List of abbreviations

LSP: Long-term stationary phase;
*E. coli*: *Escherichia coli*;
GLC+ (late-log): *E. coli* liquid cell culture in late-log phase;
GLC+ (ST): *E. coli* liquid cell culture in short-term stationary phase;
GLC−: *E. coli* liquid cell culture in minimal medium without glucose.
LSPE: *E. coli* liquid cell culture exposed to a supernatant of a long-term stationary phase cell culture;
GO term: Gene-Ontology-term.

## Additional files

### Additional file 1; File format: .pdf; Title: Figure S1

Description: Temporal changes in the viability of the *E. coli* cells after transfer to short-term stationary phase (GLC+ (ST), green), LSP supernatant (LSPE, blue), and fresh glucose-deficient medium (GLC−, magenta). An inset shows the growth curve of the cells from the late-log (black line) to the short-term stationary phase (green line). The sample preparation was performed according to the procedure explained in Fig. 1A. The *E. coli* cells were firstly cultured in fresh M63 medium with glucose for 20 h (we set this time point as 0 h in the figure) and further incubated to the short-term stationary phase or transferred to LSPE or GLC−. The number of viable cells was estimated by the CFUs. Error bars indicate standard deviations of biological triplicates.

### Additional file 2; File format: .pdf; Title: Figure S2

Description: *K-*means clustering of the samples by PC1. *K-*means clustering was performed following PCA of all samples (panel A, corresponding to Fig. 2B) or the non-growing group (panel B, corresponding to Fig. 2C) using a first principal component. Shaded areas correspond to the clusters, and the included plots are the members of each cluster. Here, a maximum number of clusters was the number of conditions in each analysis (*K* = 4 (A), *K* = 3 (B)). We changed the number of initial clusters from 1 to 17 and chose the most optimal clustering results (giving the smallest total within-cluster sum of squares) among them.

### Additional file 3; File format: .pdf; Title: Figure S3

Description: Interrelation of PCA scores and expression levels of extracted genes. The transcription levels of the extracted genes were plotted against the PCA scores. The levels of the expression in the members of PC1 top groups show a positive correlation with PC1 scores, whereas the expression levels of the genes in the bottom groups show a negative correlation with PCA scores. Each plot was colored the same as in the case of Fig. 2B.

### Additional file 4; File format: .pdf; Title: Figure S4

Description: Average mRNA signals in the extracted GO terms. Box plots of average log10 scaled mRNA signals of all members in “transport (GO:0006810),” “biosynthetic process (GO:0009058),” and “ATP synthesis coupled electron transport (GO:0042773),” which largely contribute to PC1 scores in Fig. 2B and C. Error bars indicate the standard deviation of biological replicates. Letters above each plot indicate significance groups according to Tukey’s HSD test (p < 0.05).

### Additional file 5; File format: .xls; Title: Data S1

Description: Lists of gene members in GO terms used in this study. We downloaded the original data from http://www.biocyc.org and used it with slight modifications.

### Additional file 6; File format: .xls; Title: Data S2

Description: A list of all screened GO terms, which includes a significantly larger number of highly contributing genes to PC1 in Fig. 2B and C (p-value < 0.01, hypergeometric test). For each GO term, we also showed the screened gene members, their average values of factor loading (correlation coefficient for the gene expression levels and PC scores), and p-value for the involvement of the highly contributing genes.

### Additional file 7; File format: .xls; Title: Data S3

Description: All GO terms included in “biosynthetic process (GO:0009058),” “transport (GO:0006810),” “macromolecule biosynthetic process (GO:0009059),” and “response to stress (GO:0006950)” as child classes or their parts. Those GO terms were used in the calculation of the total mRNA signals in Fig. 3B and Fig. S4 (Additional file 4).

### Additional file 8; File format: .xls; Title: Data S4

Description: All extracted GO terms significantly differentially expressed between LSPE and GLC+ (ST) (FDR q-value < 0.05, Gene Set Enrichment Analysis). Enrichment Score (ES), Normalized Enrichment Score (NES), Nominal p-value (NOM. p. val.), False Discovery Rate (FDR) are shown in the data.

### Additional file 9; File format: .xls; Title: Data S5

All extracted GO terms significantly differentially expressed between high cell density (10^9^ cells/mL) and low cell density (10^7^ cells/mL) conditions in LSPE (FDR q-value < 0.03, Gene Set Enrichment Analysis). Enrichment Score (ES), Normalized Enrichment Score (NES), Nominal p-value (NOM. p. val.), False Discovery Rate (FDR. q. val.).

**Fig. S1.**
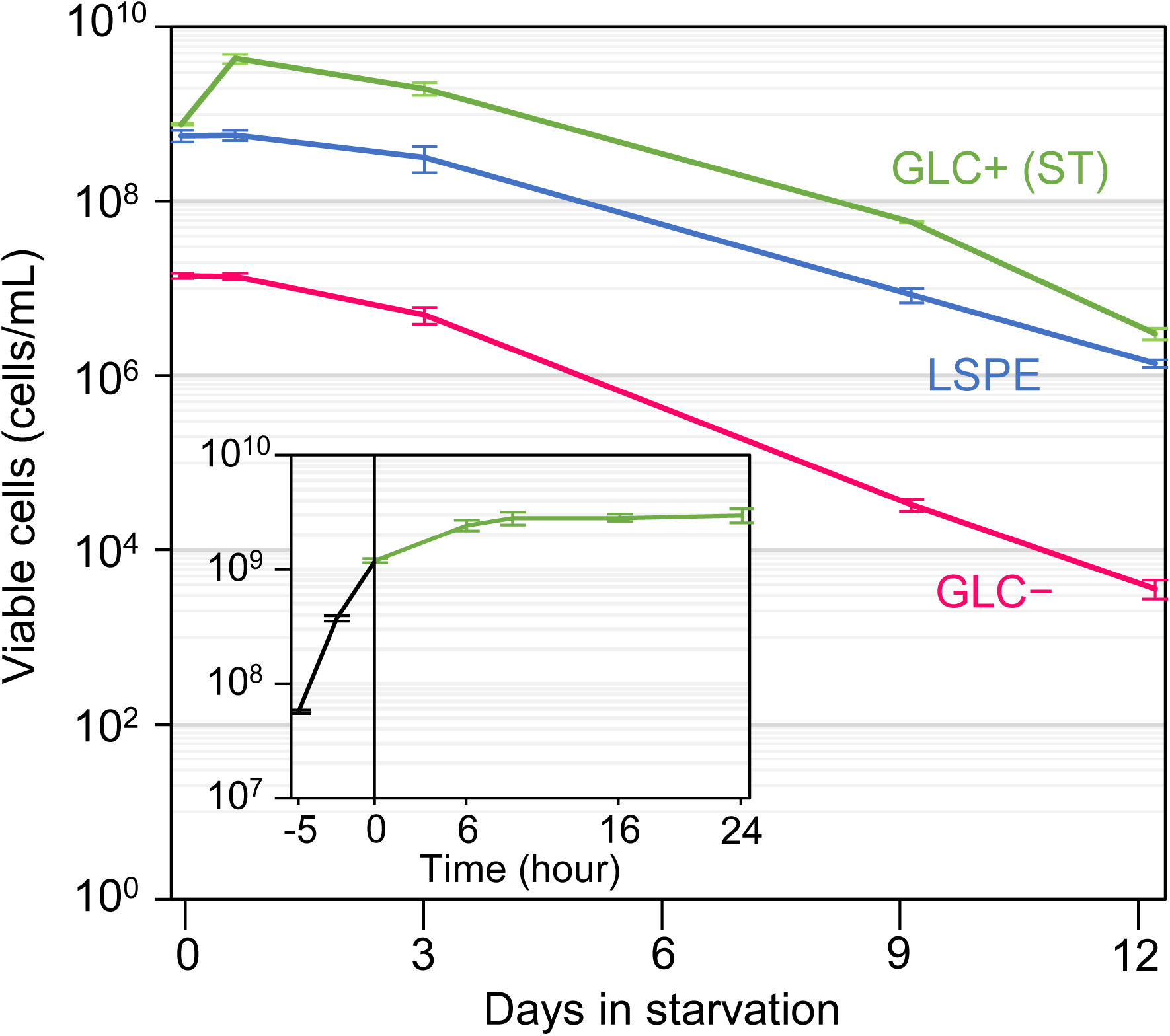

**Fig. S2.**
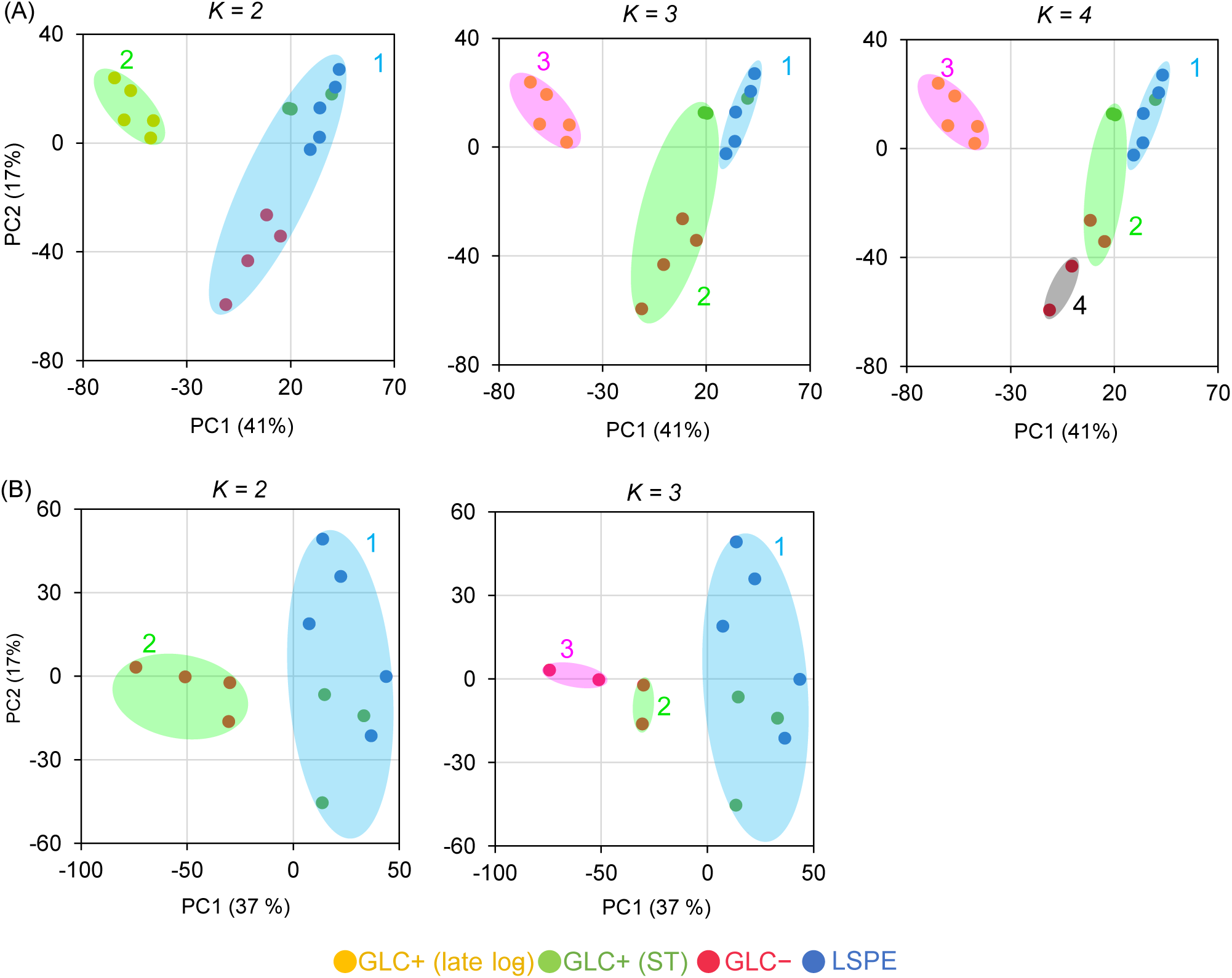

**Fig. S3.**
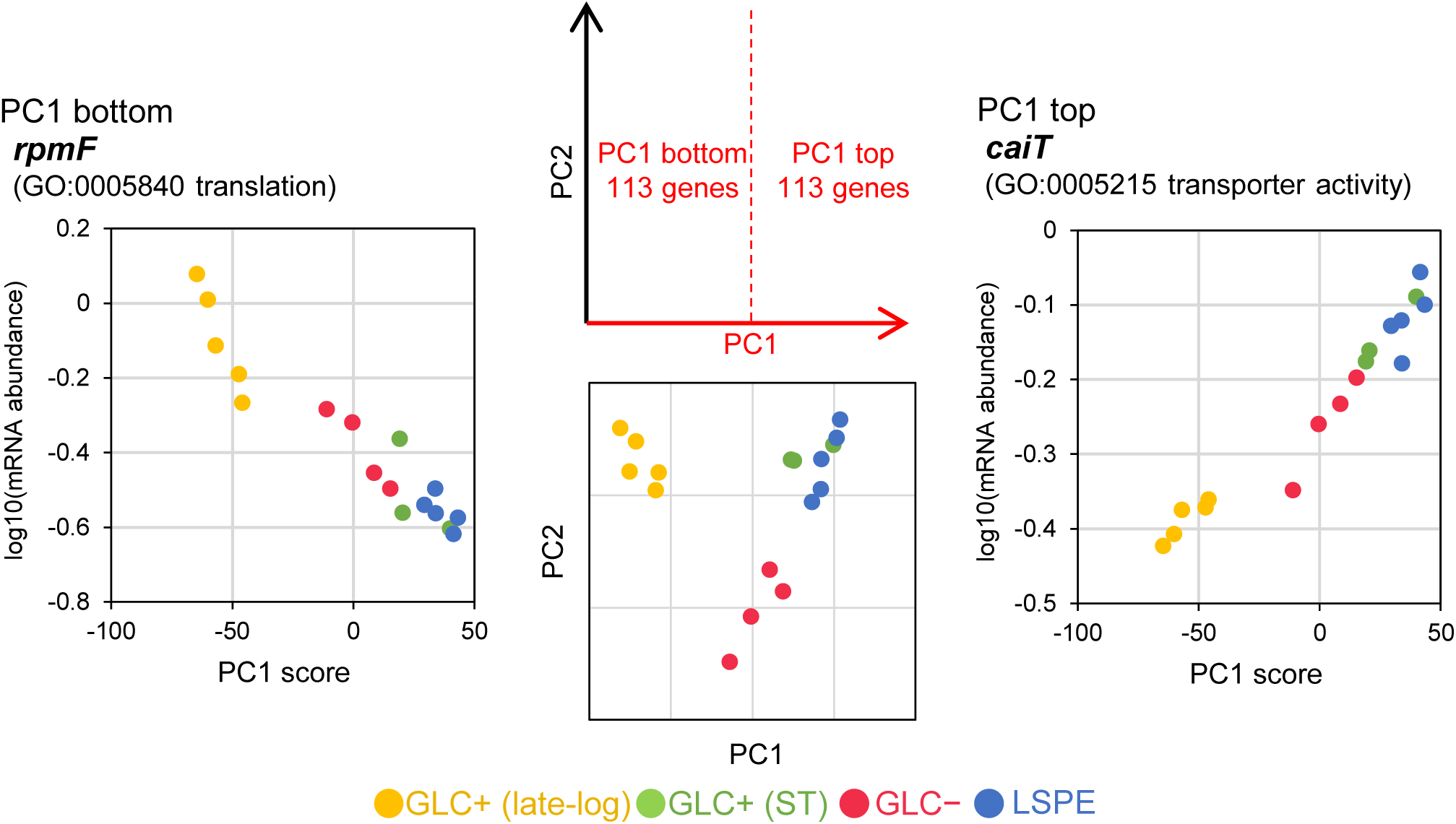

**Fig. S4.**
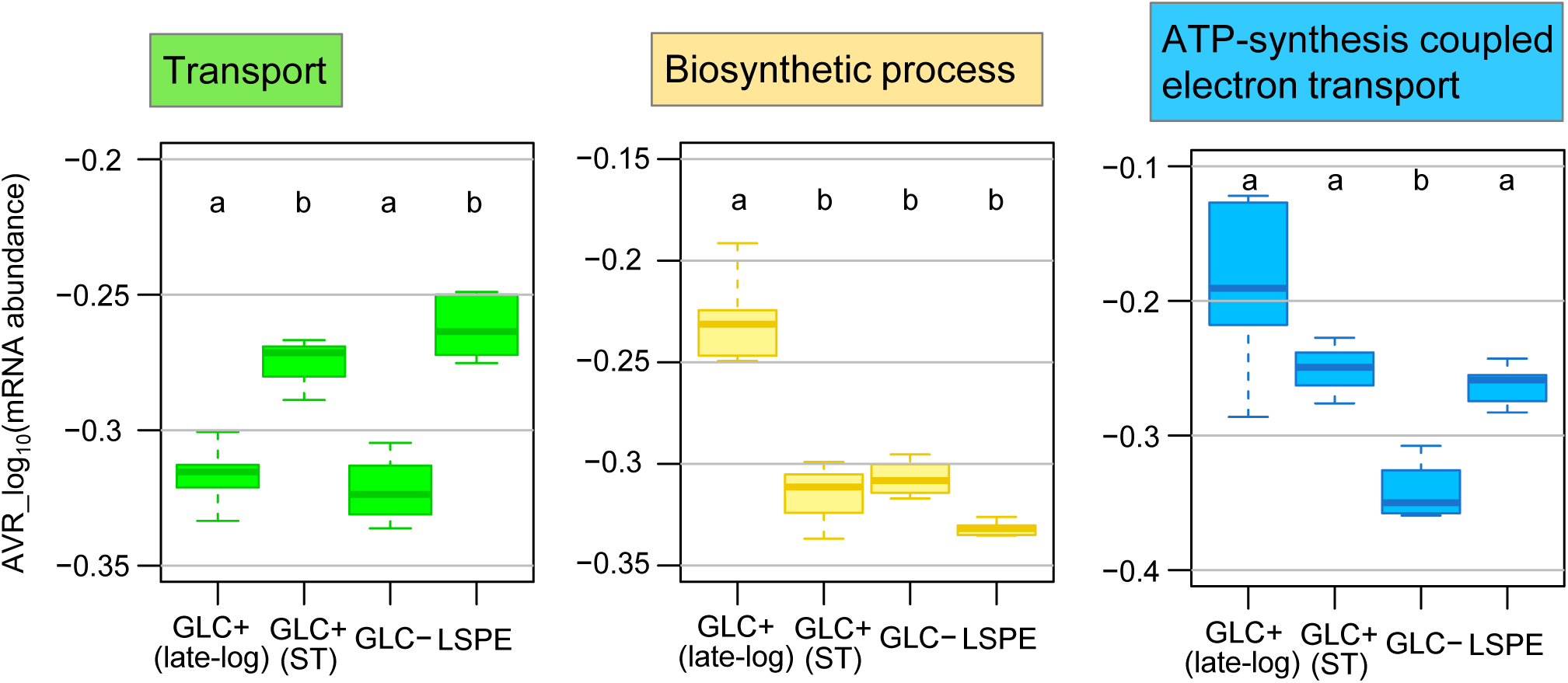

## Notes

### Competing Interest Statement

The authors have declared no competing interest.

